## supplementary results and text for "Learning low-dimensional generalizable natural features from retina using a U-net"

### Suppelmentary Information

#### A Methods

##### A.1 Finding reliable cells from a retinal population.

In our analysis, we defined reliable cells as those that fire more than 8 spikes per trial. Because each trial (movie presentation) lasts 20s, this corresponds to a mean firing rate of 0.4 spike/s. In Fig 1B, we showed a histogram of mean spikes we obtained from the entire 93-cell retinal population. We found 47 reliable cells. We averaged each cell's spiking activity over all presentations of the same movie to generate the peri-stimulus time histogram (PSTH) for that movie (e.g. fish, leaf, water). For a given 500ms window of retinal activity, we generated a unique sample by randomly dropping out 2 cells (without permutation) that fire during the specific 500ms segment. The encoder-decoder uses the resulting 45-cell retinal activity to reconstruct a movie frame 100ms beyond the specific 500ms window. For each movie, we trained our encoder-decoder to reconstruct 400 frames, using 100 unique 45-cell samples for each frame. We then tested the encoder-decoder on the first 100 frames of the 400-frame segment. We obtained another 100 unique samples of retinal activity for these test frames. These testing samples are hold-out samples that do not exist in the training dataset.

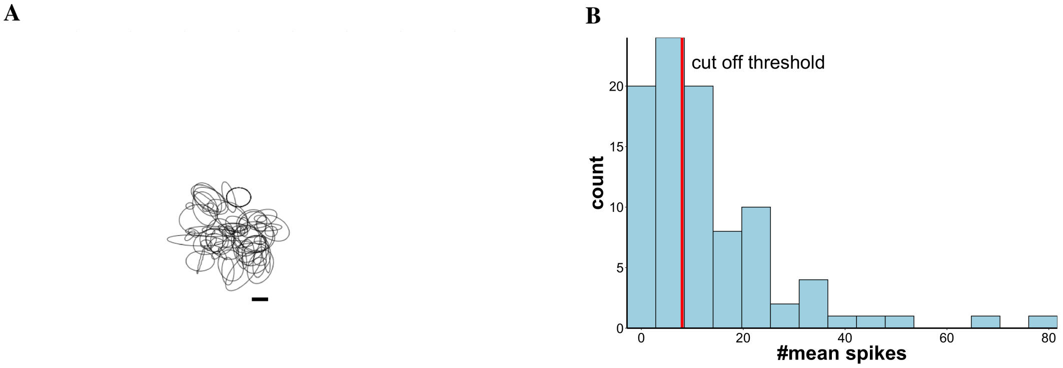


**Fig 1: A) Spatial receptive fields (RF) of all 93 retinal ganglion cells. An individual RF is a 2D Gaussian fitted to the maximum firing rate over time of a specific cell. In this retinal population, 88% are fast-off cells. B) Finding reliable cells from a retinal population. The histogram above shows the mean spikes a cell may have during a single, 20s-long trial, averaged by all trials. The red line shows the threshold by which we decide whether a cell is reliable (mean spikes > threshold) or unreliable (mean spikes < threshold). We also calculated the reliability measure used in [1] and we obtained similar results.**

##### A.2 Intrinsic Dimension Estimation

The estimation of intrinsic dimension is an active research topic. There are many estimators available. In Fig 1C of the main paper, we used the maximal likelihood based, K-nearest neighbor estimator first proposed in [2]. Using high-resolution images with known intrinsic dimension, the experiments in [3] have shown that this estimator consistently yields more accurate estimates compared to other popular methods. In our application, we estimated the intrinsic dimension with a wide range of $k$ ($k \in (5, 25)$). It is the same range used in [3]. For retinal activity, we empirically observed that these intrinsic dimension estimates converge to 2.7 when $k >= 18$. This is the "ID(PSTH)" we reported in Fig 1C. Similar to retinal activity, we also reported the converged intrinsic dimension estimate for latent activations. For example, we observed that when $k >= 11$, the intrinsic dimension estimates for a 10-D latent activation converge to $(4.0, 4.1)$ and this is what we showed in Fig 1C as well.

##### A.3 U-net

U-net was developed for biomedical image segmentation [4]. Its architecture is based on a fully convolutional network. Typical feedforward convolutional neural networks contain contracting layers only. These contracting layers form a cascade of convolution and pooling layers within which the pooling layers downsample the input. U-net is an encoder-decoder. It concatenates a contractive feedforward network (encoder) with another convolutional network of expansive layers (decoder). This gives the overall network a U-shape. U-net also appends the activations from intermediate convolutional layers of its encoder component to its decoder. These are the so-called skip connections. These skip connections are copies of the input being represented within the feature space of different intermediate convolution layers in the encoder.

*Modifications of the original U-net in our analysis:*

- We used ResNet18[5] pretrained with ImageNet[6] as the encoder, and generated an expansive decoding architecture by mirroring the encoder. There is no residual connection in the decoder component.
- The skip connections within a U-net essentially perform autoencoding. Observing that retinal activity is low-dimensional, we modified the original skip connections to become variational sampling layers [7]. The dimensionality of these variational sampling layers are parameters we varied in our analysis. To simplify training, all variational sampling layers share the same dimensionality.

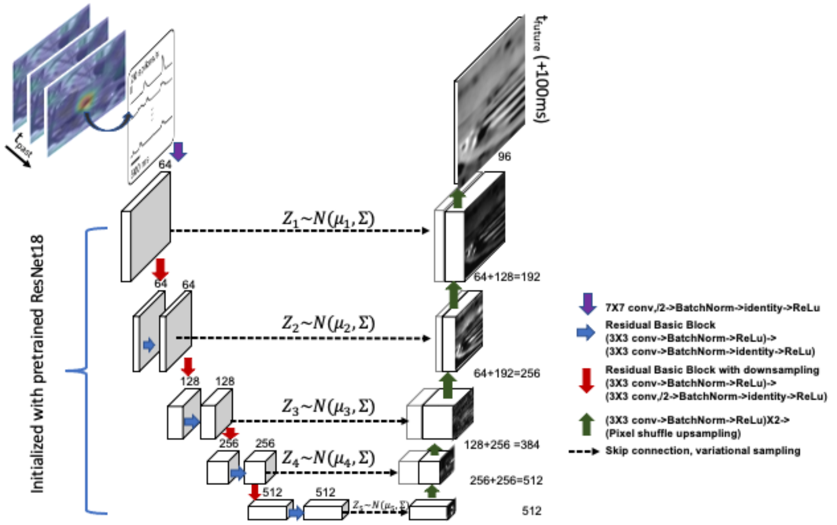


**Fig 2: The detailed U-net Diagram. This diagram corresponds to the modified U-net we used in our analysis. The encoder shares the same architecture as the ResNet18. The decoder mirrors the feedforward architecture of the encoder. The skip connections here are variational sampling layers. Each of them learns a separate latent space. To simplify training, we used the same dimensionality across all latent spaces. We also held the marginal variance $\Sigma$ fixed. Combining all skip connections, the U-net used through most of the paper has a 10D latent space in each of the 5 skip connection variational autoencoding layers, resulting in an overall 50D latent space that decodes future movie frames from retinal activity.**

##### A.4 Conversion of time-series for PSTH into a unique image

We converted all time series of mean firing rates into their respective Gramian angular fields (GAF) [8]. The Gramian angular field (GAF) represents an 1D time series with a 2D polar coordinate system. All elements in a GAF image are the trigonometric sum (i.e., superposition of directions) between different time intervals. For example, the pixel at position $(i, j)$ shows the $\cos(r_i + r_j)$ for a PSTH sequence $\{r_1, \cdots, r_i, \cdots, r_j, \cdots, r_n\}$. This method has been shown to successfully capture high fluctuations in financial time series [9]. In our analysis, we first normalized all mean firing rates into $(0, 1)$ per 500ms segment. We then computed the $\cos(r_i + r_j)$ to fill up off-diagonal $(i, j)$ pixels with $i \neq j$. Lastly, we filled the diagonal $(i, i)$ pixels of a GAF image with the raw, unnormalized mean firing rates.

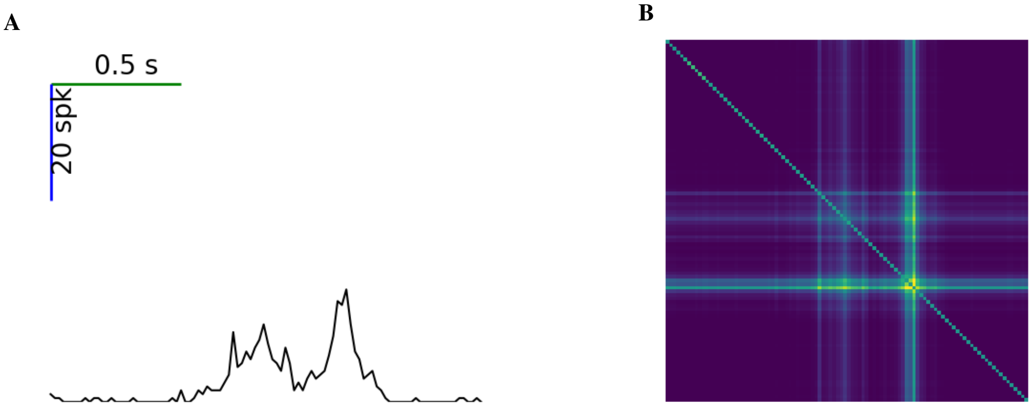


**Fig 3: A) Mean firing rates $r_{1,\cdots,n}$ of a sample neuron. B) The Gramian angular field corresponding to A). Values at the diagonal pixels are raw mean firing rates, the same as A). Values at the off-diagonal pixels $(i, j)$ are $\cos(r_i, r_j)$ after $r_{1,\cdots,n}$ is normalized to $(0, 1)$, respectively.**

##### A.5 Training details

We used a cyclical learning rate schedule [10, 11] to speed up our training. The maximum learning rate is set to $1e-2$. The specific learning rate per epoch is determined by the loss automatically [12]. We observed that 20 epochs (2-3 hours on an NVIDIA Tesla K80 GPU) are generally enough to train an entire encoder-decoder. We split the 20 epochs into two phases. In the first phase, we only trained the decoder and froze the weights of the encoder component (initialized by the pretrained ResNet18). In the second phase, we unfroze the entire model to train both the encoder and the decoder together. To prevent overfitting, we reduced the maximum learning rate if the improvement between epochs is less than 0.1. We also performed early stopping if the improvement is less than 0.1 between two consecutive epochs.

##### A.6 Represent a natural image with features from the pretrained VGG19

Natural images contain high level features beyond their pixels. Pretrained CNNs capture these high-level features in their convolutional layers. By feeding a natural image into a pretrained CNN, one may obtain a representation that contains a whole suite of features from narrow to broad spatial scales [13]. VGG19 [14] is one of the most widely used pretrained CNN. Previous works have shown that convolutional layers of the pertained VGG19 provide features that can predict both neural responses and human eye movements [15, 16, 17]. A specific convolutional layer, the 'conv5_4', has been shown to contain features that can predict video saliency in humans [18]. To perform hierarchical clustering on frames, we used feature activations from this convolutional layer ('conv5_4') as a generic representation for both movie frames and optic flow frames.

##### A.7 Obtain distributions for static/dynamic motifs of a natural movie based on hierarchical clustering

For all three movies, we created the clustering hierarchy of their 100 test frames and their optic flow frames separately using agglomerative hierarchical clustering [19]. This clustering algorithm first creates a matrix of pairwise distance between all frames. Because each image is represented by their features from the pretrained VGG19, this pairwise distance corresponds to how two images activate the pretrained VGG19 differently. There are 512 features in the 'conv5_4' layer of the VGG19. Using feature activation of a 64X64 image from these 512 features, we performed principal component analysis and found that the first 10 principal components of these activations explain $\sim$95% of variance in the data. This observation holds for all three movies and their respective optic flow frames. Therefore, we calculated the pairwise distance between frames as the Euclidean distance using these 10 principal components. The clustering algorithm takes a bottom-up approach to build the clustering hierarchy from the matrix of pairwise distance. It starts with each frame as an individual cluster. It merges pairs of frames, or pairs of clusters as it moves up the hierarchy. The choice of merging two clusters greedily minimizes the total within-cluster variance (the ward's criterion [20]). The algorithm takes 99 merging steps to cluster 100 frames. It terminates when the hierarchy reaches 1 cluster.

Every time the algorithm merges two clusters into one, it outputs a distance between the two clusters that are being merged. This distance shows how similar/dissimilar these two clusters are. Because this agglomerative hierarchical clustering is greedy, this distance only increases as the algorithm moves up the hierarchy. We generated discrete distributions of static or dynamic motifs by thresholding clustering hierarchies based on this distance. Such thresholding results in all clusters at the level right below the specific threshold being treated as discrete states of a probability distribution for the 100 test frames. They are shown in Fig 4A,5A,6A for static movie frames and Fig 4B,5B,6B for dynamic optic flow frames. Using a low threshold, one may obtain a distribution with many clusters. Each of them may contain a few frames only. This distribution will have many states and a high entropy. Using a high threshold, many frames are grouped into one cluster and the corresponding distribution will have a low entropy. For all hierarchical clustering, we observed that this clustering distance grows slowly in the beginning, and increases exponentially towards the end. We performed a grid search on possible thresholds for each movie. Our goal is to construct $Y_{dynamic}$, $Y_{static}$ such that the mutual information $I(Y_{static}, Y_{dynamic})$ is small. "Small" here is defined such that $I(Y_{static}, Y_{dynamic})$ may only take up to 50% of the entropy in both $H(Y_{dynamic})$ and $H(Y_{static})$. In Fig 4D,Fig 5D and Fig 6D, we reported a "redundancy ratio"( i.e., $I(Y_{static}, Y_{dynamic})/H(Y_{static})$ and $I(Y_{static}, Y_{dynamic})/H(Y_{dynamic})$) for possible thresholds. Among all thresholds that yield a small $I(Y_{static}, Y_{dynamic})$ (below the cut-off redundancy ratio = 0.5), we prefer a low threshold such that the resulting $Y_{joint}$ may retain more information within $H(time)$. For two movies (fish and leaf), we identified a threshold within the slow growing phase (highlighted between two dashed lines in Fig 4C and Fig 5C). For the water movie, we only found a high threshold with which the $I(Y_{static}, Y_{dynamic})$ is small.

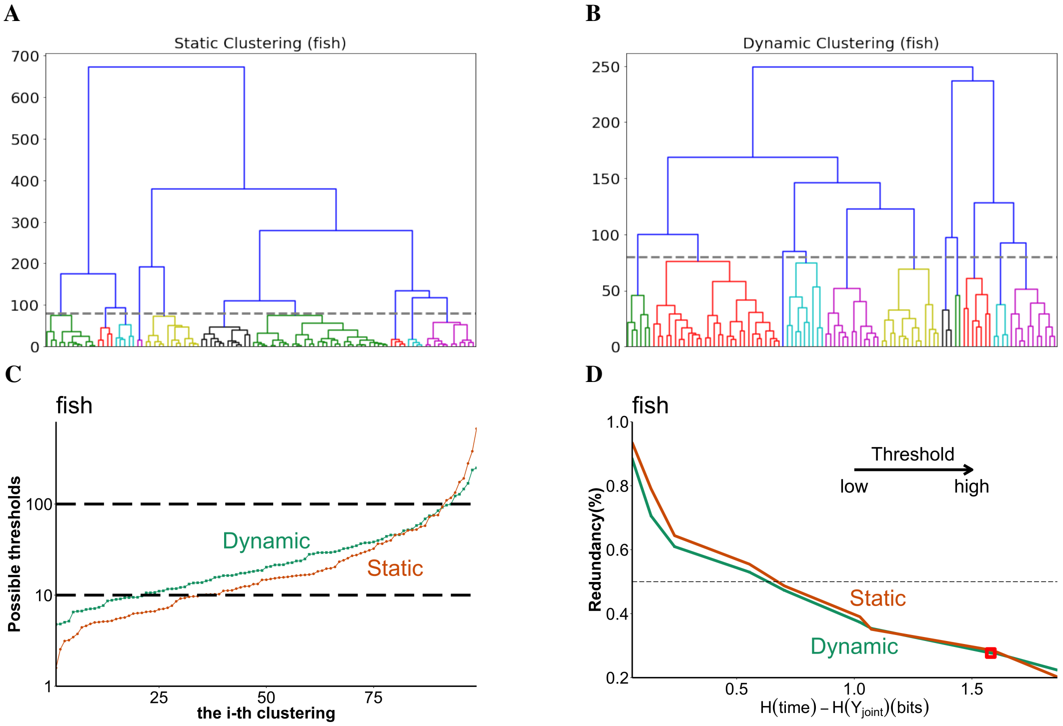


**Fig 4: A) The agglomerative hierarchical clustering using static features of the test segment (0-100th) of the fish movie. The threshold that generates $Y_{static}$ in Fig 5 of the main paper is shown as the gray dashed line. All clusters are shown with different colors. Because each cluster groups multiple frames together (10 clusters for 100 frames), this probability distribution is also a coarse-grained distribution of time within the 100-frame movie segment. B) The same as A), but using optic flow frames (dynamic features) of the test segment. C) The distance of all 99 merging steps in logarithmic scale as the algorithm builds up its clustering hierarchy. This distance grows slowly in the beginning and exponentially towards the end. We performed a grid search within the slow-growing phase (highlighted by dashed lines) to find the thresholds used in A) and B). D) As the threshold increases (moves up the clustering hierarchy), redundancy ratios of both static/dynamic features decrease. Meanwhile, $Y_{joint}$ also loses information about time. The $I(Y_{static}, Y_{dynamic})$ is small if both redundancy ratios $I(Y_{static}, Y_{dynamic})/Y_{static}$ and $I(Y_{static}, Y_{dynamic})/Y_{static}$ are less than 0.5 (dashed line). The red square highlights the $Y_{joint}$ (its entropy and redundancy ratios) we used in Fig 5 of the main paper. The results we reported in Fig 5 of the main paper hold for all $Y_{joint}$ with redundancy ratios<=0.5.**

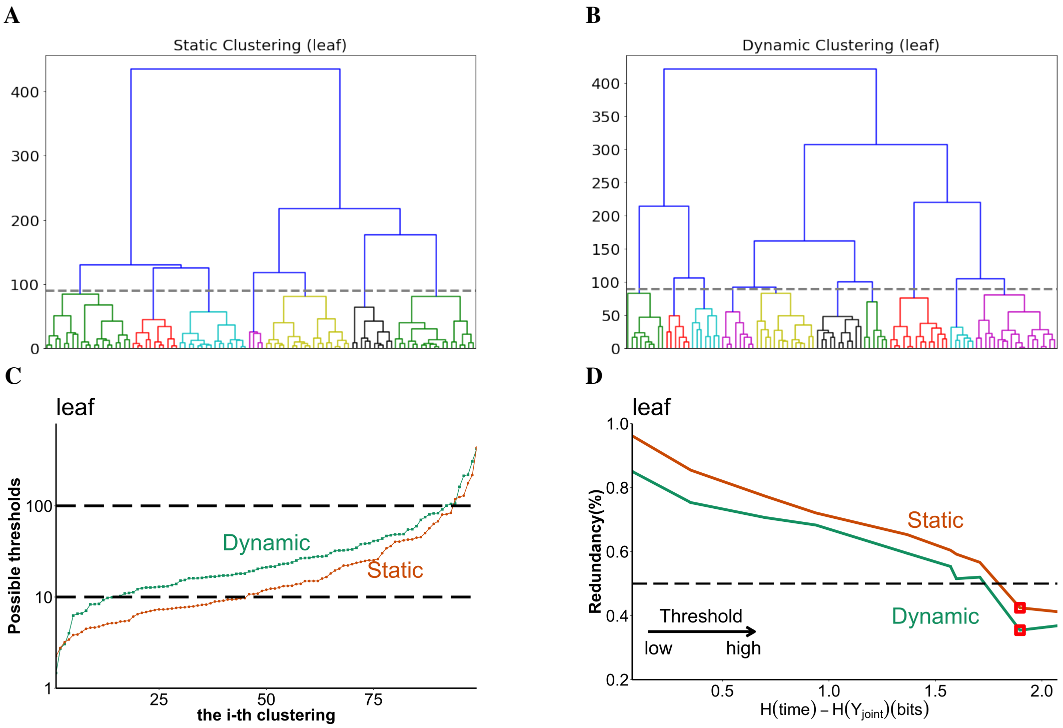


**Fig 5: The same as Fig 4, but for the leaf movie**

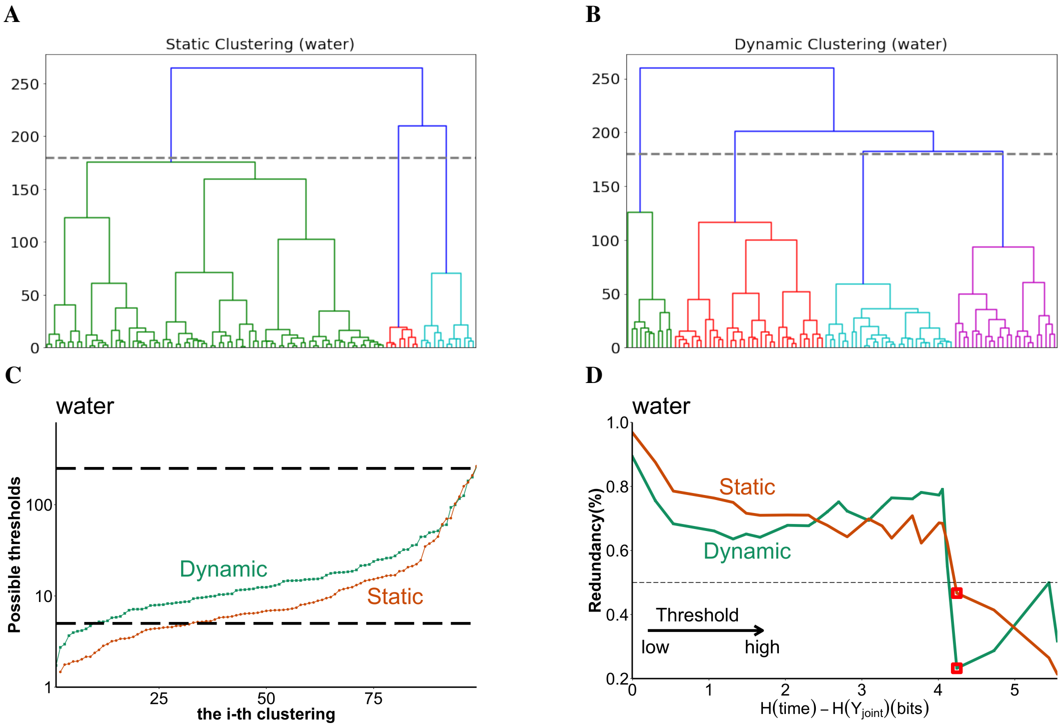


**Fig 6: The same as Fig 4 and Fig 5, but for the water movie. A) The same as Fig 5A and Fig 4A, but for static frames of the water movie. B) The same as Fig 5B and Fig 4B, but for optic flow frames of the water movie. C) Similar to Fig 5C and Fig 4C, but we expanded the grid search of possible thresholds to cover most of the distance span used by hierarchical clustering. D) We could only obtain a small $I(Y_{static}; Y_{dynamic})$ with a high threshold. The corresponding $H(Y_{joint})$ contains 2.4 bits only, about 37% of $H(time)$.**

#### B Additional Results

##### B.1 Encoder-decoders trained with the other two movies (leaf and water) also learn a generalizable encoding of time in the natural scene up to its full entropy

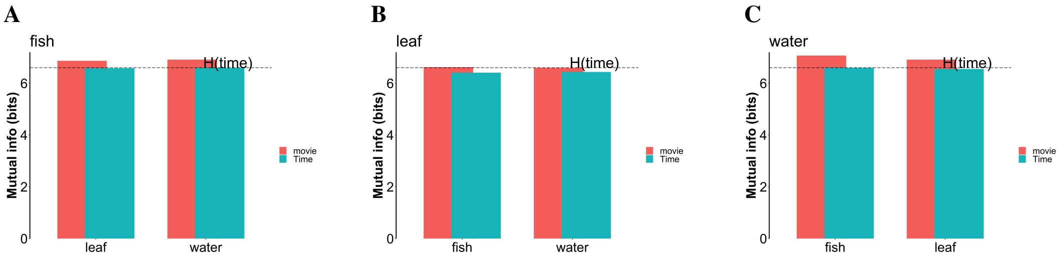


**Fig 7: A) The same as Fig 3B of the main paper, we included it here for completeness. B) The same as A), but using an encoder-decoder trained to decode frames of the leaf movie from retinal activity. Red bars show $I(Z_{leaf}; Z_{fish})$ and $I(Z_{leaf}; Z_{water})$. Cyan bars show the fraction within $I(Z_{leaf}; Z_{fish})$ and $I(Z_{leaf}; Z_{water})$ that is about time. C) The same as A) and B), but using an encoder-decoder trained to decode frames of the water movie from retinal activity.**

| | Water(5d) | Water(10d) | Leaf(5d) | Leaf(10d) |
|---|---|---|---|---|
| Fish | 78.2% | 96.9% | 72.2% | 98.0% |
| Leaf | 79.5% | 97.8% | 84.4% | 99.1% |
| Water | 84.0% | 97.4% | 71.7% | 97.9% |

**Table 1: Latent representations from encoder-decoder trained on any one movie can decode time in all three movies. Here we showed two configurations (latent dimension =5 or 10, respectively).**

##### B.2 Synergistic features in the other two (leaf and water) movies

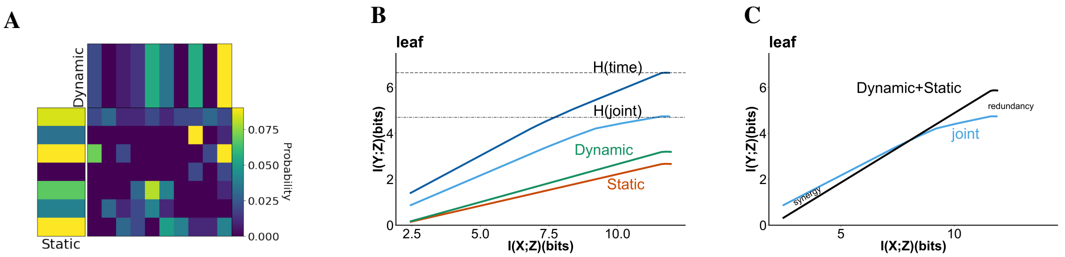


**Fig 8: The same as Fig 5 of the main paper, but for the leaf movie. A) Joint distribution of static and dynamic features. This joint distribution includes 71% of $H(time)$. B) The information plane for leaf data. Dark blue: the information curve for encoding time; Light blue: the information curve for encoding the joint distribution combining static and dynamic features; Red/Green: information curves for separated static (red) and dynamic (green) features. C) Blue: the information curve for encoding the joint distribution, the same as B); Black: the sum of information curves from Dynamic(Red)+Static(Green). There is a synergistic region between the information curve for the joint and the sum. This synergistic region is slightly smaller compared to what we observed in the fish movie (Fig 5C of the main paper).**

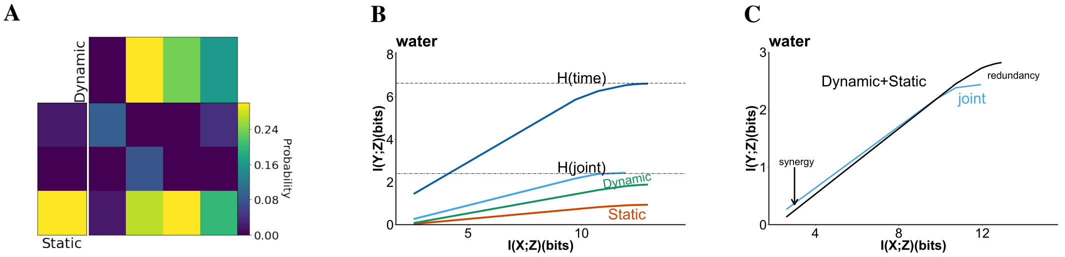


**Fig 9: The same as Fig 5 of the main paper, but for the water movie. A) Joint distribution of static and dynamic features. Because of the high threshold shown in Fig 6D, this joint distribution contains less number of discrete states compared to the joint distributions for the other two movies. B) The information plane for water data. Dark blue: the information curve for encoding time; Light blue: the information curve for encoding the joint distribution combining static and dynamic features; Red/Green: information curves for separated static (red) and dynamic (green) features. C) Blue: the information curve for encoding the joint distribution, the same as B); Black: the sum of information curves from Dynamic(Red)+Static(Green). The synergistic region is visible, but smaller compared to what we observed using the other two (leaf and fish) movies.**

##### B.3 Videos

Here is the detailed list of 11 videos we included in the supplementary material:

1. Natural movie segments: (fish, leaf, water)movie.mp4. These movies are segments from the Chicago motion Database [21]. All movies are 400-frame long and a frame rate of 60fps. After training, we tested our encoder-decoder with 10,000 hold-out retinal activity for the 100 frames from the beginning of these segments.
2. Test segments and their reconstructions: (fish, leaf, water)_truthvspred.mp4. We attached these movies to show the performance of three encoder-decoders (each trained for a specific movie). These movies play the target test movie segments and their respective reconstruction side by side.
3. Optic flow of the 100-frame test segments from all three movies: (fish, leaf, water)flow100frames.mp4. These optic flow frames are computed as observed motion between two consecutive frames using a pretrained FlowNet2 [22].
4. Visualization of features within a trained U-net: we included two movies to show features in the encoder-decoder trained for the fish movie. The feature_motionbackground.mp4 contains the top 5% activated features (16 out of 384) from the decoding layer we showed

| | Instantaneous | Raw PSTH | Shuffled PSTH | Isomap | 10D-PCA | 50D-PCA |
|---|---|---|---|---|---|---|
| Fish | 3.9% | 99% | 60.2% | 14.9% | 55.9% | 97.2% |
| Leaf | 4.5% | 70% | 5.6% | 7.0% | 2.5% | 59.2% |
| Water | 4.7% | 99% | 65.9% | 16.6% | 32.1% | 85.2% |

**Table 2: Simple linear decoders cannot reproduce the decoding performance we obtained from our variational U-net.**

in Fig 2 of the main paper. We used a different colormap to highlight the object motion and background features we showed in Fig 2. The feature_output.mp4 contains the top 5% activated features (4 out of 96) from the decoding layer that outputs reconstructed movie frames. The lower right feature of feature_output.mp4 shows fish movement only, similar to the object motion feature in feature_motionbackground.mp4.

##### B.4 Simple visualizations and decoding methods

We include below simple visualizations and decoding performance with simple models per the reviewers' suggestions. These visualizations show that "time in natural scene" is not a feature that is trivially encoded by the retinal population (mean PSTH), nor its generalization can be trivially observed by correlating frames between movies (pairwise frame-to-frame distance). We then use simple methods to decode this "time in natural scene" from raw PSTHs, and their dimensionality reductions. All these calculations show that finding a generalizable low dimensional feature space for all three natural movies is nontrivial.

###### B.4.1 Simple visualization I: mean PSTH of the retinal population.

Here, we show the mean PSTH of the entire retinal population (the gray region is the standard error per time bin). They do not show a clear trend that correlates with time (e.g., neurons that fire more at the onset of the movie and slowly decay to 0 by the end.) We also show the 400ms PSTH patterns from 5 example neurons responding to the fish movie. Their individual spiking patterns also do not correlate in any simple way with time since movie onset.

###### B.4.2 Simple visualization II: Frame-to-frame distance between different movies

Here we show the pairwise frame-to-frame distance between different movies. This follows the reviewer's advice to "take frames from the fish movie and compute the pairwise distance to all frames in another movie." We hypothesize that if some trivial visual features of one movie can encode time of a different movie, then there may be a correlation between frame-to-frame distance and time when both frames correspond to the same time in both movies. We use the same frame-to-frame distance in Fig 4 of the main paper. Similar to mean PSTHs, we do not observe a clear trend that correlates with time.

###### B.4.3 Simple visualization III: 2D dimensionality reduction of the latent space

Although a linear decoder can decode time from a 10D latent representation, time is not encoded by any easily discernible trivial aspect of the latent space. When we convert the 10D latent activations into 2D with Isomap, we do not observe any obvious correlation between how latent activations change through time and time itself. (Filenames are fish/leaf/water_latent.gif)

###### B.4.4 Linear decoder performance using raw PSTH's and their simple dimensionality reductions

We show here the performance (percentage correct) of 3 different linear decoders using raw PSTH's to decode time. A linear decoder trained on shuffled PSTH's (shuffled neuron identity per input) shows inferior performance compared to one trained on raw PSTH's. This means that decoding time does not come from trivial gross changes in spiking statistics as time passes during the movie. All

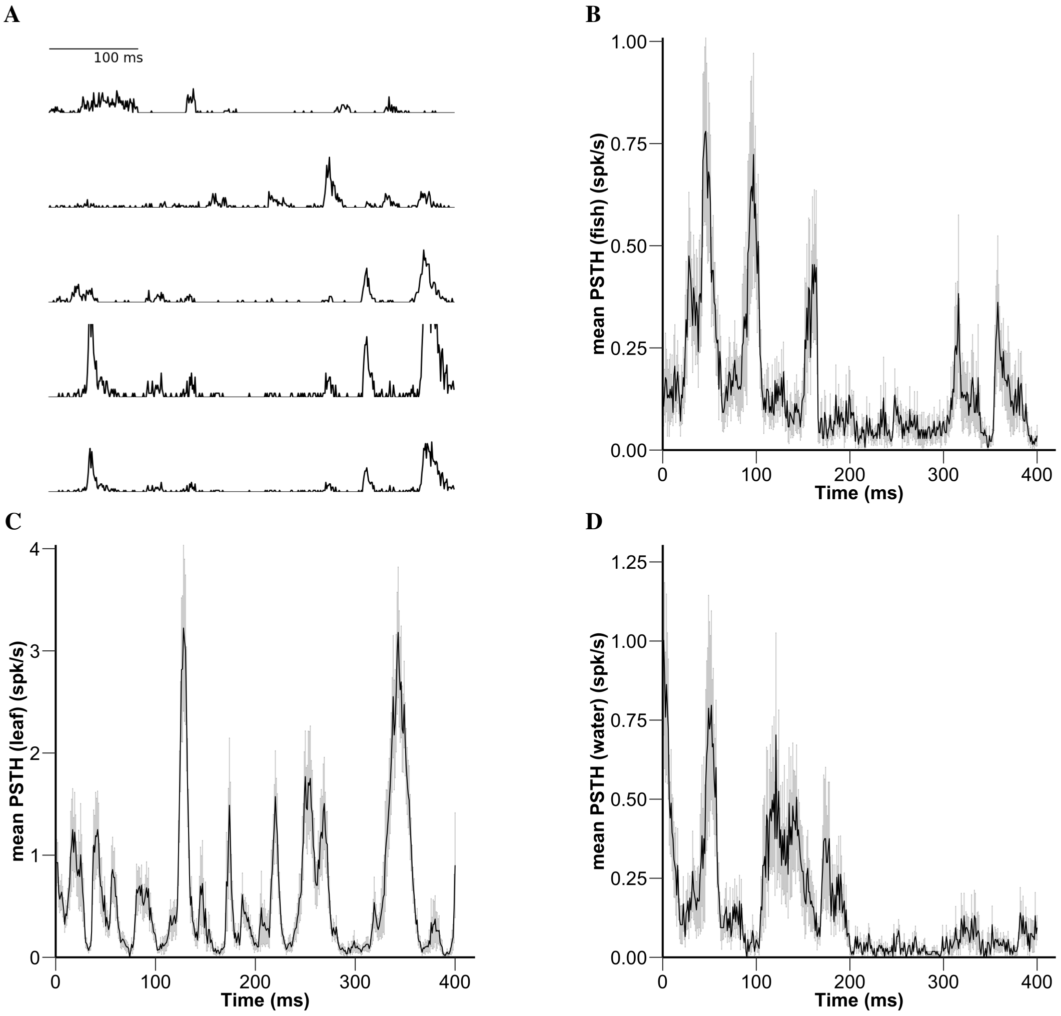


**Fig 10**

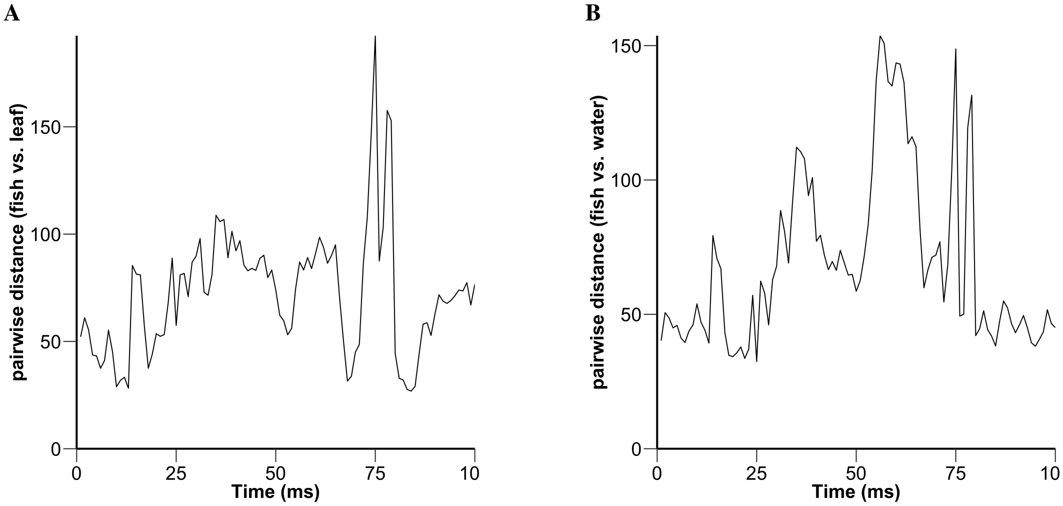


**Fig 11**

these raw PSTH's are high-dimensional: 45 cells X 30 time bins (17ms each) = 1350 dimensions. A linear decoder trained on these high-dimensional raw PSTH's works well for two movies (fish and water), but much worse for the leaf movie. It is possible that there are different nonlinear components in the retina code responding to different movies. This shows a clear advantage of U-net: a linear decoder trained on the U-net's latent representation with only 10 dimensions can decode time in the natural scene with 99

Next we investigate whether simple dimensionality reduction methods can also find low-dimensional, generalizable features for the time in the natural scene. We use two off-the-shelf dimensionality reduction methods to obtain low-dimensional representations of raw PSTH's: One is ISOMAP. It is a nonlinear dimensionality reduction method. We choose ISOMAP over another popular option (tSNE) because ISOMAP allows us to train an embedding space using the fish movie onto which we can project the other two movies. The other is PCA. It is linear. We choose d=10 because the U-net-learned latent representation achieves a 99% decoding performance with d=10. We find that neither methods show decoding performance that are comparable to what we achieve using the latent representation from a trained U-net. It is surprising how bad the ISOMAP is. Using PCA, the decoding performance varies significantly across different movies. Echoing from the decoding result using raw PSTH's, this suggests that retinal activity responding to these natural movies have complex and diverse structures. This diversity makes it challenging to discover the generalizable features across different movies.

Table 2 shows that the shuffled PSTH's perform better than using the 10D PCA in the fish movie. However, only 31.7% of the test samples are decoded correctly by both the shuffled PSTH and the 10D-PCA. Additionally, the decoding performances for the fish movie are 9.3% and 17.8% if we project the shuffled PSTH onto the subspace of 10D-PCA or 50D-PCA, respectively. As a result, there are few linear similarities between the features obtained via PCA and shuffled PSTH's. Meanwhile, we also found that, even with the 50D-PCA, linear decoders are unable to generalize as well as the 10-D latent representation learned by the U-net. Our variational U-net performs substantial nonlinear transformation to learn a low dimensional feature space applicable to all three natural movies.

### References


[1] Lane McIntosh, Niru Maheswaranathan, Aran Nayebi, Surya Ganguli, and Stephen Baccus. Deep learning models of the retinal response to natural scenes. In D. Lee, M. Sugiyama, U. Luxburg, I. Guyon, and R. Garnett, editors, *Advances in Neural Information Processing Systems*, volume 29. Curran Associates, Inc., 2016.

[2] David J.C. MacKay and Zoubin Ghahramani. Comments on 'maximum likelihood estimation of intrinsic dimension' by e. levina and p. bickel (2004). http://www.inference.org.uk/mackay/dimension/.

[3] Phillip Pope, Chen Zhu, Ahmed Abdelkader, Micah Goldblum, and Tom Goldstein. The intrinsic dimension of images and its impact on learning. April 2021.

[4] Olaf Ronneberger, Philipp Fischer, and Thomas Brox. U-net: Convolutional networks for biomedical image segmentation. May 2015.

[5] Kaiming He, Xiangyu Zhang, Shaoqing Ren, and Jian Sun. Deep residual learning for image recognition. In *2016 IEEE Conference on Computer Vision and Pattern Recognition (CVPR)*. IEEE, jun 2016.

[6] Alex Krizhevsky, Ilya Sutskever, and Geoffrey E Hinton. Imagenet classification with deep convolutional neural networks. In F. Pereira, C.J. Burges, L. Bottou, and K.Q. Weinberger, editors, *Advances in Neural Information Processing Systems*, volume 25. Curran Associates, Inc., 2012.

[7] Diederik P Kingma and Max Welling. Auto-encoding variational bayes. December 2013.

[8] Zhiguang Wang and Tim Oates. Imaging time-series to improve classification and imputation. In *Proceedings of the 24th International Conference on Artificial Intelligence*, IJCAI'15, page 3939–3945, Buenos Aires, Argentina, 2015. AAAI Press.

[9] Silvio Barra, Salvatore Mario Carta, Andrea Corriga, Alessandro Sebastian Podda, and Diego Reforgiato Recupero. Deep learning and time series-to-image encoding for financial forecasting. *IEEE/CAA Journal of Automatica Sinica*, 7(3):683–692, may 2020.

[10] Leslie N. Smith. Cyclical learning rates for training neural networks. June 2015.

[11] Leslie N. Smith and Nicholay Topin. Super-convergence: Very fast training of neural networks using large learning rates. August 2017.

[12] Jeremy Howard et al. fastai. `https://github.com/fastai/fastai`, 2018.

[13] Joris Guerin, Stephane Thiery, Eric Nyiri, Olivier Gibaru, and Byron Boots. Combining pretrained cnn feature extractors to enhance clustering of complex natural images. *Guerin, J., Thiery, S., Nyiri, E., Gibaru, O., Boots, B. (2021). Combining pretrained CNN feature extractors to enhance clustering of complex natural images. Neurocomputing, 423, 551-571*, January 2021.

[14] Karen Simonyan and Andrew Zisserman. Very deep convolutional networks for large-scale image recognition. September 2014.

[15] Matthias Kümmerer, Lucas Theis, and Matthias Bethge. Deep gaze i: Boosting saliency prediction with feature maps trained on imagenet. November 2014.

[16] Matthias Kummerer, Thomas S.A. Wallis, Leon A. Gatys, and Matthias Bethge. Understanding low- and high-level contributions to fixation prediction. In *2017 IEEE International Conference on Computer Vision (ICCV)*. IEEE, oct 2017.

[17] Santiago A. Cadena, George H. Denfield, Edgar Y. Walker, Leon A. Gatys, Andreas S. Tolias, Matthias Bethge, and Alexander S. Ecker. Deep convolutional models improve predictions of macaque v1 responses to natural images. *PLOS Computational Biology*, 15(4):e1006897, apr 2019.

[18] Matthias Tangemann, Matthias Kümmerer, Thomas S.A. Wallis, and Matthias Bethge. Measuring the importance of temporal features in video saliency. *Journal of Vision*, 20(11):1061, oct 2020.

[19] F. Pedregosa, G. Varoquaux, A. Gramfort, V. Michel, B. Thirion, O. Grisel, M. Blondel, P. Prettenhofer, R. Weiss, V. Dubourg, J. Vanderplas, A. Passos, D. Cournapeau, M. Brucher, M. Perrot, and E. Duchesnay. Scikit-learn: Machine learning in Python. *Journal of Machine Learning Research*, 12:2825–2830, 2011.

[20] Joe H. Ward Jr. Hierarchical grouping to optimize an objective function. *Journal of the American Statistical Association*, 58(301):236–244, 1963.

[21] Jared Salisbury and Stephanie Palmer. Chicago motion database. https://cmd.rcc.uchicago.edu/.

[22] Eddy Ilg, Nikolaus Mayer, Tonmoy Saikia, Margret Keuper, Alexey Dosovitskiy, and Thomas Brox. Flownet 2.0: Evolution of optical flow estimation with deep networks. December 2016.
